# Learning predictive cognitive maps with spiking neurons during behaviour and replays

**DOI:** 10.1101/2021.08.16.456545

**Authors:** Jacopo Bono, Sara Zannone, Victor Pedrosa, Claudia Clopath

## Abstract

We describe a framework where a biologically plausible spiking neural network mimicking hippocampal layers learns a cognitive map known as the successor representation. We show analytically how, on the algorithmic level, the learning follows the TD(*λ*) algorithm, which emerges from the underlying spike-timing dependent plasticity rule. We then analyze the implications of this framework, uncovering how behavioural activity and experience replays can play complementary roles when learning the representation of the environment, how we can learn relations over behavioural timescales with synaptic plasticity acting on the range of milliseconds, and how the learned representation can be flexibly encoded by allowing state-dependent delay discounting through neuromodulation and altered firing rates.

## 1. Introduction

Mid twentieth century, Tolman proposed the concept of cognitive maps [Tolman, 1948]. These maps are abstract mental models of an environment which are helpful when learning tasks and in decision making. Since the discovery of hippocampal place cells, cells that are activated only in specific locations of an environment, it is believed that the hippocampus can provide the substrate to encode such cognitive maps [O’Keefe and Dostrovsky, 1971; O’Keefe, J.; Nadel, 1978].

While these place cells offer striking evidence in favour of cognitive maps, it is not clear what representation is actually learned by the hippocampus and how this information is exploited when solving and learning tasks. Recently, it was proposed that the hippocampus computes a cognitive map containing predictive information, called the successor representation (SR). Theoretically, this SR framework has some very interesting properties, such as efficient learning, simple computation of the values of states, fast relearning when the rewards change and flexible decision making [Dayan, 1993; Stachenfeld et al., 2014, 2017; Russek et al., 2017; Momennejad et al., 2017]. Furthermore, the SR reproduces some important experimental observations. Firstly, the firing fields of hippocampal place cells are affected by the strategy used by the animal to navigate the environment (known as the *policy* in machine learning), as well as by changes in the environment [Mehta et al., 2000; Stachenfeld et al., 2017]. Secondly, reward revaluation — the ability to recompute the values of the states when rewards change — would be more effective than transition revaluation – the ability to recompute the values of states when the environment changes [Russek et al., 2017; Momennejad et al., 2017].

In this work, we study how this predictive representation could be implemented by biologically plausible neural networks with spike-timing dependent synaptic plasticity (STDP). While STDP is the underlying learning rule, we show that on an algorithmic level the learning is equivalent to TD(*λ*), a well studied and powerful algorithm known from reinforcement learning [Sutton and Barto, 1998].

Our proposed framework has some interesting implications. In particular, we will discuss how it unifies rate coding and temporal coding, and how we can learn relationships between states that are seconds apart while using STDP timescales in the millisecond range. Subsequently, we will analyze the delay discounting parameter *γ* in our model. We will discuss how the relationship between *γ* and time allows us to consider time as a continuous variable, therefore we don’t need to discretize time as is usual in reinforcement learning. Moreover, the delay-discounting in our model depends hyperbolically on time but exponentially on state transitions. We will discuss how the *γ* parameter can be modulated by neuronal firing rates and neuromodulation, allowing state-dependent discounting and in turn enabling richer information in the SR such as salient states, landmarks, reward locations, etc. Finally, we will investigate the perks of using replays when learning the cognitive map. Following properties of TD(*λ*), we show how we can achieve both low bias and low variance by using replays, translating to both quicker initial learning and convergence to lower error. We will then discuss how we can use replays to learn *offline*. In this way, policies can be refined without the need for actual exploration.

Our framework allows us to make predictions about the roles of behavioural learning and replay-like activity and how they can be exploited in representation learning. Furthermore, we uncover a relation between STDP and a higher level learning algorithm. In this way, our work spans the three levels of analysis proposed by Marr [Marr, 2010]. On the implementational level, our model consists of a feedforward network of excitatory neurons with biologically plausible spike-timing dependent plasticity. On the algorithmic level, we show that our model learns the successor representation using the TD(*λ*) algorithm. On the computational theory level, our model tackles representation learning using cognitive maps.

## 2. Results

Cognitive maps are internal models of an environment, helping animals to learn, plan and make decisions during task completion. The hippocampus has long been thought to provide the substrate for learning such cognitive maps, and recent evidence points towards a specific type of representation learned by the hippocampus, the successor representation (SR) [Stachenfeld et al., 2017].

### 2.1 The Successor Representation

In this section, we will give an overview of the successor representation and its properties, especially geared towards neuroscientists. Readers already familiar with this representation may safely move to the next section.

To understand the concept of successor representation (SR), we can consider a spatial environment — such as a maze — while an animal explores this environment. In this setting, the SR can be understood as how likely it is for the animal to visit a future location starting from its current position. We further assume the maze to be formed out of a discrete number of states. Then, the SR can be more formally described by a matrix with dimension (*N*_*states*_ × *N*_*states*_), where *N*_*states*_ denotes the number of states in the environment and each entry *R*_*ij*_ of this matrix describes the expected future occupancy of a state *S*_*j*_ when the current state is *S*_*i*_. In other words, starting from *S*_*i*_, the more likely it is for the animal to reach the location associated with state *S*_*j*_ and the nearer in the future, the higher the value of *R*_*ij*_.

As a first example, we consider an animal running through a linear track. We assume the animal runs at a constant speed and always travels in the same direction — left to right (Figure 1a). We also split the track into four sections or *states, S*_1_ to *S*_4_, and the SR will be represented by a matrix with dimension (4 × 4). Since the animal always runs from left to right, there is zero probability of finding the animal at position *i* if its current position is greater than *i*. Therefore, the lower triangle of the successor matrix is equal to zero (Figure 1b). Alternatively, if the animal is currently at position *S*_1_, it will be subsequently found at positions *S*_2_, *S*_3_, and *S*_4_ with probability 1. The further away from *S*_1_, the longer it will take the animal to reach that other position. In terms of the successor matrix, we apply a discounting factor *γ* (0 < *γ* ≤ 1) for each extra “step” required by the animal to reach a respective location (Figure 1b).

**Figure 1.**
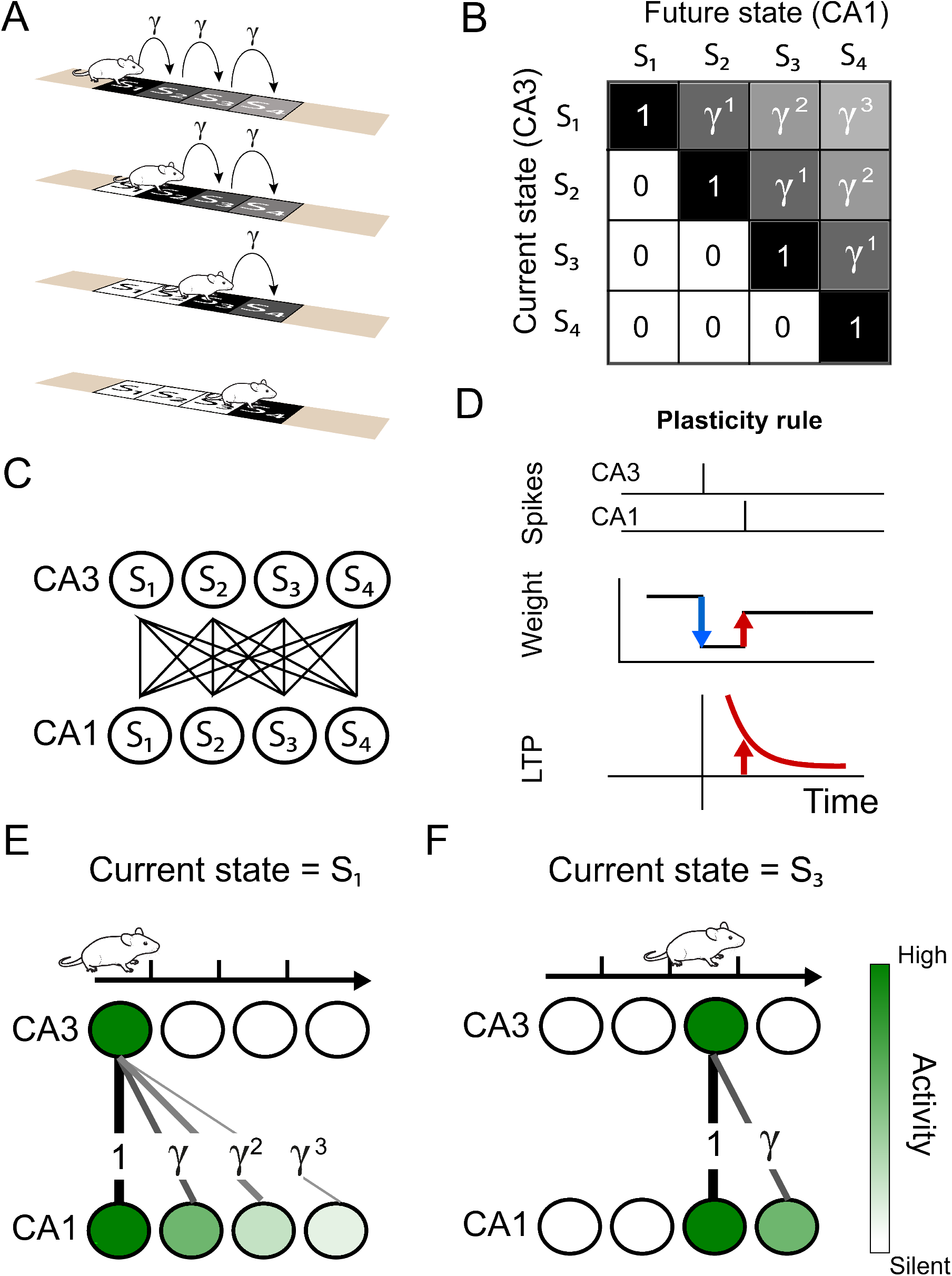
Successor representation and neuronal network. (A) Our simple example environment consists of a linear track with 4 states (S_1_ to S_4_) and the animal always moves from left to right — i.e. one epoch consists of starting in S_1_ and ending in S_4_. (B) The successor matrix corresponding to the task described in panel A. (C) Our neuronal network consists of a 2 layers with all-to-all feedforward connections. The presynaptic layer mimics hippocampal CA3 and the postsynaptic layer mimics CA1. (D) The synaptic plasticity rule consists of a depression term and a potentiation term. The depression term is dependent on the synaptic weight and presynaptic spikes (blue). The potentiation term depends on the timing between a pre- and postsynaptic spike pair (red), following an exponentially decaying plasticity window (bottom). (E-F) Schematics illustrating some of the results of our model. (E) Our spiking model learns the top row of the successor representation (panel B) in the weights between the first CA3 place cell and the CA1 cells. (F) Our spiking model learns the third row of successor representation (panel B) in the weights between the third CA3 place cell and the CA1 cells.

Note that the representation learned by the SR is dependent not only on the structure of the environment, but also on the policy — or strategy — used by animal to explore the environment. In this first example, the animal applied a deterministic policy (always running from left to right), but the SR can also be learned for stochastic policies. Furthermore, the SR is a multi-step representation, in the sense that it stores predictive information of multiple steps ahead. Because of this predictive information, the SR retains some important advantages usually attributed to model-based learning such as sample-efficient re-learning when reward location is changed [Gershman, 2018]. In model-based learning, differently from the SR, the single-step transition probabilities between all states of the environment are learnt, which can be computationally expensive. The SR, however, can be efficiently learned using model-free methods and allows us to easily compute values for each state, which in turn can guide the policy [Dayan, 1993]. This position between model-based and model-free methods makes the SR framework very powerful, and its similarities with hippocampal neuronal dynamics has led to increased attention from the neuroscience community. Finally, in our examples above we considered an environment made up by a discrete number of states. This framework can be generalised to a continuous environment represented by a discrete number of place cells.

### 2.2 Learning the successor representation in biologically plausible networks

We propose a model of the hippocampus that is able to learn the successor representation. We consider a feedforward network comprising of two layers. Similar to [Mehta et al., 2000], we assume that the presynaptic layer represents the hippocampal CA3 region and is all-to-all connected to a postsynaptic layer - representing the CA1 network (Figure 1c). The synaptic connections from CA3 to CA1 are plastic such that the weight changes follow a spike-timing dependent plasticity (STDP) rule consisting of two terms: a weight-dependent depression term for presynaptic spikes and a potentiation term for pre-post spike pairs (Figure 1d).

For simplicity, we assume that the animal spends a fixed time *T* in each state. During this time, a constant activation current is delivered to the CA3 neuron encoding the current location and, after a delay, to the corresponding CA1 place cell (see Methods). On top of these fixed and location-dependent activations, the CA3 neurons can activate neurons in CA1 through the synaptic connections. In other words, the CA3 neurons are activated according to the current location of the animal, while the CA1 neurons have a similar location-dependent activity combined with activity caused by presynaptic neurons. The constant currents delivered directly to CA3 and CA1 neurons can be thought of as location-dependent currents from entorhinal cortex. These activations subsequently trigger plasticity at the synapses, and we can show analytically that, using the spike-timing dependent plasticity rule discussed above, the SR is learned in the synaptic weights (Figure 1e,f, and see supplementary methods).

Moreover, we find that, on an algorithmic level, our weight updates are equivalent to a learning algorithm known as TD(*λ*). TD(*λ*) is a powerful and well-known algorithm in reinforcement learning that itself unifies two families of learning algorithms known as Monte Carlo (MC) learning and temporal difference (TD) learning [Sutton and Barto, 1998]. It is characterized by the parameter *λ*, which controls how close the algorithm is to Monte Carlo or TD learning. The extreme cases TD(1), equivalent to MC, and TD(0), or simply TD, have different strengths and weaknesses, as we will discuss in more detail in the next sections.

Importantly, from our analytical analysis (see supplementary methods), we find that the *λ* parameter depends on the behavioural parameter T (the time an animal spends in a state). We find that, the larger the time T, the smaller the value of *λ* and vice-versa. In other words, when the animal moves through the trajectory on behavioural time-scales (large T compared to the synaptic plasticity time-scales *τ*_LTP_), the network is learning the SR with TD(*λ* ∼ 0). For quick sequential activities (T → 0), akin to hippocampal replays, the network is learning the SR with TD(*λ* ∼ 1). As we will discuss below, this framework therefore elegantly combines learning based on rate coding as well as temporal coding. Furthermore, from our model follows the prediction that replays can also be used for learning purposes and that they are algorithmically equivalent to MC, whereas during behaviour, the hippocampal learning algorithm is equivalent to TD. This strategy of using replays to learn is in line with recent experimental and theoretical observations (see [Momennejad, 2020] for a review).

To validate our analytical results, we use again a linear track with a deterministic policy. Using our spiking model with either rate-code activity on behavioural time-scales (Figure 2a top) or temporal-code activity similar to replays (Figure 2b top), we show that the synaptic weights across trials match the evolution of the TD(*λ*) algorithm closely (Figure 2a,b middle). After convergence, the final weight matrix indeed corresponds to the SR matrix simulated through TD(*λ*) (Figure 2a,b bottom).

**Figure 2.**
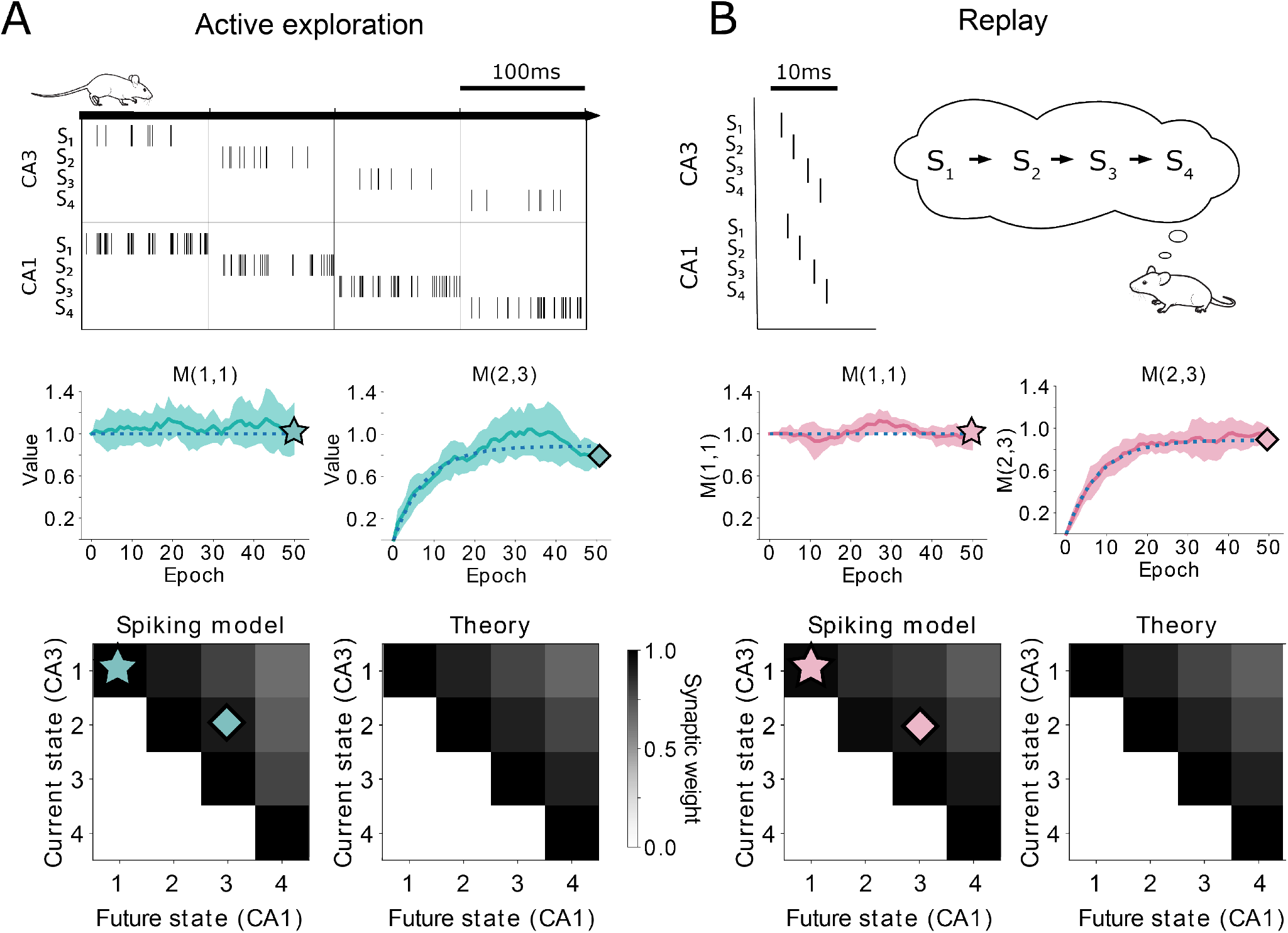
Comparison between TD(*λ*) and our spiking model. (A-top) Learning during behaviour corresponds to TD(*λ* ≈ 0). States are traversed on timescales larger than the plasticity timescales and place cells use a rate-code. (A-middle) Comparison of the learning over epochs for two synaptic connections (full line denotes the mean over ten random seeds, shaded area denotes one standard deviation) with the theoretical learning curve of TD(*λ*) (dotted line). (A-bottom) Final successor matrix learned by the spiking model (left) and the theoretical TD(*λ*) algorithm (right). Star and diamond symbols denote the corresponding weights shown in the middle row. (B-top) Learning during replays corresponds to TD(*λ* ≈ 1). States are traversed on timescales similar to the plasticity timescales and place cells use a temporally precise code. (B-middle and bottom) Analogous to panel A middle and bottom.

In summary, we showed how the network can learn the SR using a spiking neural model. We analytically showed how the learning algorithm is equivalent to TD(*λ*), and confirmed this using numerical simulations. We derived a relationship between the abstract parameter *λ* and the timescale T representing the animal’s behaviour — and in turn the neuronal spiking — allowing us to unify rate and temporal coding within one framework. Furthermore, we predict a role for hippocampal replays in learning the SR using an algorithm equivalent to Monte Carlo.

### 2.3 Learning over behavioural time-scales using STDP

An important observation in our framework is that the SR can be learned *using the same underlying STDP rule* over time-scales ranging from replays up to behaviour. One can now wonder how it is possible to learn relationships between events that are seconds apart during awake behaviour, without any explicit error encoding signal typically used by the TD algorithm, and while the STDP rule is characterised by millisecond time-scales (Figure 3a).

**Figure 3.**
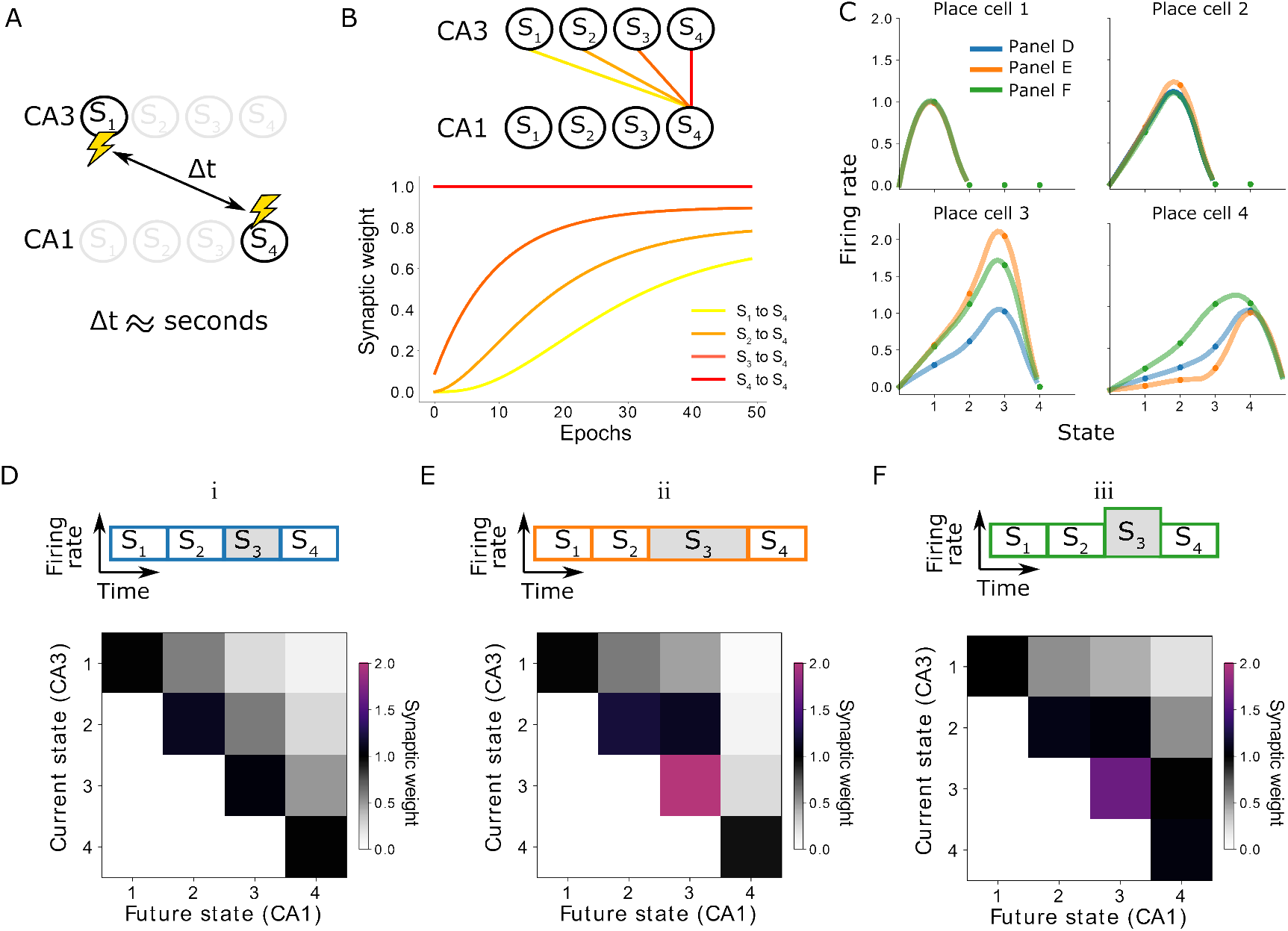
Learning on behavioural timescales and state-dependent discounting. (A) In our model, the network can learn relationships between neurons that are active seconds apart, while the plasticity rule acts on a millisecond timescale. (B) Due to transitions between subsequent states, each synaptic weight update depends on the weight from the subsequent CA3 neuron to the same CA1 neuron. In other words, the change of a synaptic weight depends on the weight below it in the successor matrix. The top panel visualizes how weights depend on others in our linear track example, where each lighter colour depends on the darker neighbour. The bottom panel shows the learning of over 50 epochs. Notice the lighter traces converge more slowly, due to their dependence on the darker traces. (C) Place fields of the place cells in the linear track — each place cell corresponding to a column of the successor matrix. Activities of each place cell when the animal is in each of the four states (dots) are interpolated (lines). Three variations are considered: (i) the time spent in each state and the CA3 firing rates are constant (blue and panel D); (ii) the time spent in state 3 is doubled (orange and panel E); (iii) the CA3 firing rate in state 3 is doubled (green and panel F). Panels E and F lead to a modified discount parameter in state 3, affecting the receptive fields of place cells 3 and 4

From a neuroscience perspective, this can be understood when considering the trajectory of the animal. Each time the animal moves from a position *S*_*j*−1_ to a position *S*_*j*_, the CA3 cell encoding the location *S*_*j*−1_ stops firing and the CA3 cell encoding the location *S*_*j*_ starts firing. Since in our example this transition is instantaneous, these cells are activating the same CA1 cells consecutively. Therefore, the change in the weight *w*_*i,j*−1_ depend on the synaptic weight of the subsequent state *w*_*i,j*_ (Figure 3b, yellow depends on orange, orange depends on red, etc). Indeed, in our example of an animal in a linear track subdivided into four locations, the weights on the diagonal, such as *w*_4,4_, are the first one to be learned, since they are learned directly. The off-diagonal weights, such as *w*_3,4_, *w*_2,4_, and *w*_1,4_, are learned consecutively more slowly as they are dependent on the subsequent synaptic weight. Eventually, weights between neurons encoding positions that are behaviorally far apart can be learnt using a learning rule on a synaptic timescale (Figure 3b).

From a reinforcement learning perspective, the TD(*λ*) algorithm relies on a property called bootstrapping. Bootstrapping means that a function is updated using current estimates of the same function, and lies at the heart of TD algorithms. The parameter *λ* in TD lambda controls the amount of bootstrapping. The case in which *λ* = 0 is equivalent to one-step TD, which fully bootstraps: at every timestep, we calculate a *prediction error*, i.e. the error between the current estimate of our SR matrix and the observed state occupancy (where the agent actually lands). We then use this prediction error to update and correct our SR estimate (see Methods).

Importantly, the prediction error in our model is not encoded by a separate mechanism in the way that dopamine is thought to do for reward prediction [Schultz et al., 1997]. Instead, the prediction error is represented locally, at the level of the synapse, through the depression and potentiation terms of our STDP rule, and the current weight encodes the current estimate of the SR. The bootstrapping in our spiking model arises due to the dependency of synaptic weights on each other, as discussed above. Notably, the prediction error updates are split into two separate synaptic changes in our model, but result in a total update equivalent to the TD(*λ*) update. We therefore do not need a vector of prediction errors as proposed in [Gardner et al., 2018; Gershman, 2018]. In fact, the synaptic potentiation in our model updates a row of the SR, while the synaptic depression updates a column.

On the other extreme, for very fast timescales such as replays, TD(1) is equivalent to online Monte Carlo learning (MC), which does not bootstrap at all. Instead, MC samples the whole trajectory and then simply takes the average of the discounted state occupancies to update the SR (see Methods). During replays, the whole trajectory falls under the plasticity window and the network can learn without bootstrapping. For all cases in between, the network partially relies on bootstrapping and we correspondingly find a *λ* between 0 and 1.

In summary, in our framework synaptic plasticity leads to the development of a successor representation in which synaptic weights can be directly linked to the successor matrix. In this framework, we can learn over behavioural timescales even though our plasticity rule acts on the scale of milliseconds, due to the bootstrapping property of TD algorithms.

### 2.4 Different discounting for space and time

In reinforcement learning, it is usual to have delay-discounting: rewards that are further away in the future are discounted compared to rewards that are in the immediate future. Intuitively, it is indeed clear that a state leading to a quick reward can be regarded as more valuable compared to a state that only leads to an equal reward in the distant future. For tasks in a tabular setting, with a discrete state space and where actions are taken in discrete turns, such as for example chess or our simple linear track discussed in section 2.1, one can simply use a multiplicative factor 0 < *γ* ≤ 1 for each state transition. In this case the discount follows an exponential dependence, where rewards that are *n* steps away are discounted by a factor of *γ*^*n*^.

In order to still use the above exponential discount when time is continuous, the usual approach is to discretize time by choosing a unit of time. However, this would imply one can never remain in a state for a fraction of this unit of time, and it is not clear how this unit would be chosen. Our framework deals naturally with continuous time, through the monotonically decreasing dependence of the discount parameter *γ* on the time an agent remains in a state, T. The dependence on T can be interpreted as an increased discounting the longer a state lasts.

In this way, instead of discounting by *γ*^*n*^ when the agent stays *n units of time* in a certain state, we would discount by *γ*(*n* · *T*). More generally, for any arbitrary time T, a discount corresponding to *γ*(*T*) will be applied. This allows the agent to act in continuous time (Figure 3c, e). Interestingly, the dependence of *γ* on T in our model is not exponential as in the tabular case. Instead, we have a hyperbolic dependence. This hyperbolic discount is well studied in psychology and neuroeconomics and appears to agree well with experimental results [Laibson, 1997; Ainslie, 2012].

The difference between a hyperbolic discount and an exponential discount lays in the fact that we will attribute a different value to the same temporal delay, depending on whether it happens sooner or later. A classic example is that, when given the choice, people tend to prefer 100 dollars today instead of 101 dollars tomorrow, while they tend to prefer 101 dollars in 31 days instead of 100 dollars in 30 days. They therefore judge the 1 day of delay differently when it happens later in time. Exponential discounting, on the other hand, always attributes the same value to the same delay no matter when it occurs.

Our model therefore elegantly combines two types of discounting: exponential when we move through space — i.e. when sequentially activating different place cells — and hyperbolic when we move through time — i.e. when we prolong the activity of the same place cell.

The discount factor *γ* also depends on other parameters such as firing rate and STDP amplitudes. This gives our model the flexibility to encode state-dependent discounting even when the trajectories and times spent in the states are the same. Such state-dependent discounting can be useful to for example encode salient locations in the environment such as landmarks or reward locations (Figure 3c, f).

### 2.5 Bias-variance trade-off

As discussed previously (section 2.2), the TD(*λ*) algorithm unifies the TD algorithm and the MC algorithm. In our framework, replay-like neuronal activations are equivalent to MC, while behavioural-like activity is equivalent to TD. In this section we will discuss how the replays and behaviour can work together when learning the cognitive map of an environment, leveraging the strengths of MC and TD.

The MC algorithm samples a full trajectory and updates the estimate of the SR with that sample. In the stochastic setting, where there is some randomness in choosing actions or transitioning states, such samples contain a lot of variance. Therefore, our MC estimate of the SR is said to have high variance as well. However, on average, our prediction of the SR is accurate, and our MC estimate of the SR is said to have low bias (Figure 4A,B).

**Figure 4.**
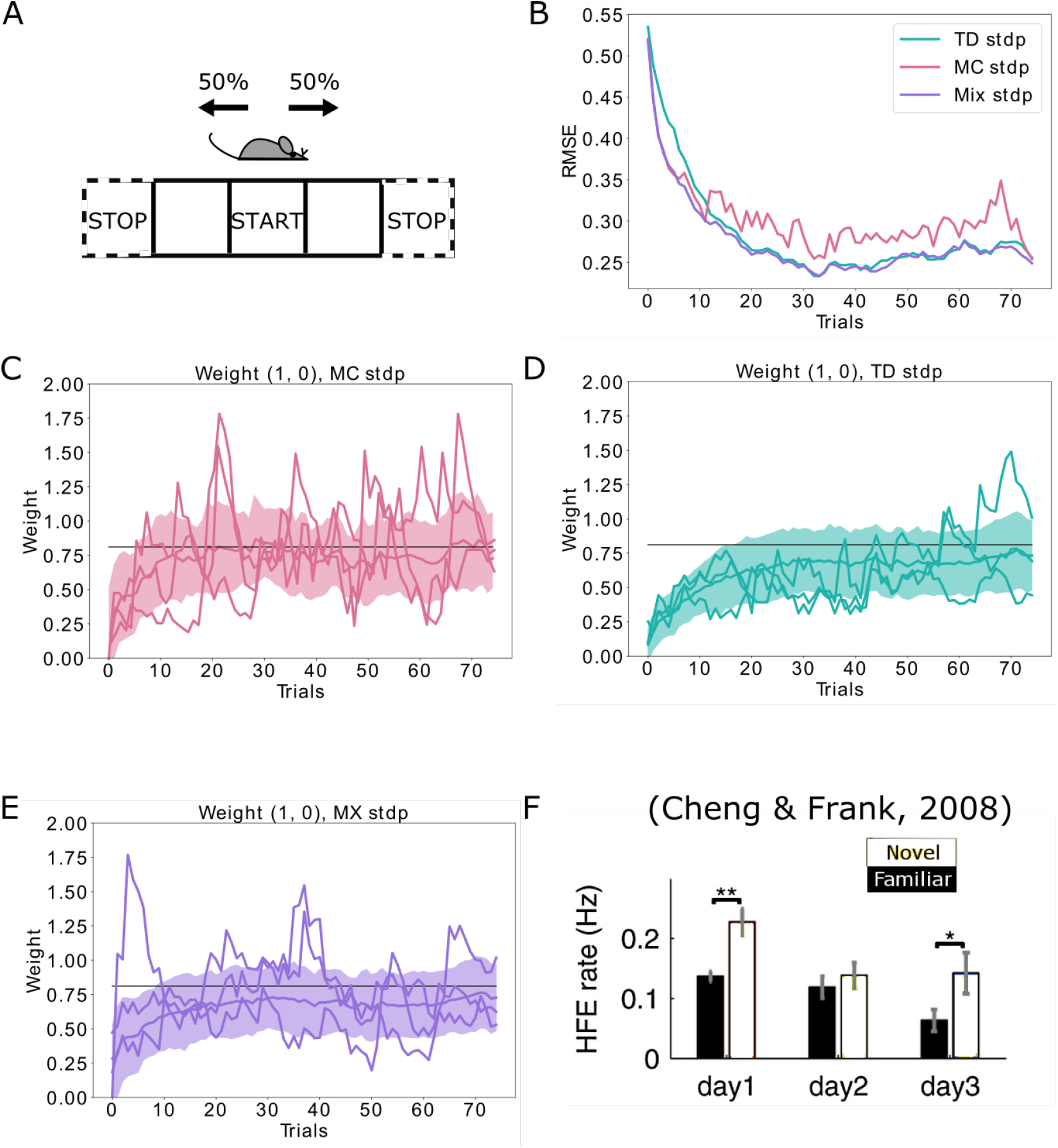
Replays can be used to control the bias-variance trade-off. (A) The agent follows a stochastic policy starting from the initial state (denoted by *START*). The probability to move to either neighbouring state is 50%. An epoch stops when reaching a terminal state (denoted with *STOP*). (B) Root mean squared error (RMSE) between the learned SR estimate and the theoretical SR matrix. The full lines are mean RMSEs over 1000 random seeds. Three cases are considered: (i) learning happens exclusively due to behavioural activity (TD STDP, green); (ii) learning happens exclusively due to replay activity (MC STDP, purple); (iii) A mixture of behavioural and replay learning, where the probabilities for replays drops off exponentially with epochs (Mix STDP, pink). The *mix* model, with a decaying number of replays learns as quickly as MC in the first epochs and converges to a low error similar to TD, benefiting both from the low bias of MC at the start and the low variance of TD at the end. (C, D, E) Representative weight changes for each of the scenarios. Full lines show various random seeds, shaded areas denote one standard deviation over 1000 random seeds. (F) More replays are observed when an animal explores a novel environment (day 1). Figure reproduced from Cheng and Frank [2008]

Unlike MC, the TD algorithm updates its estimate of the SR by comparing the current estimate of the SR and the actual state the agent transitioned to. Because of the dependence on the current estimate, this estimate will be incrementally refined with small updates. The SR estimate will in this way be lower in variance. However, by this dependence on the current estimate, we introduce a bias in the algorithm, which will be especially significant when our initial estimate of the SR is bad (Figure 4A,B).

Initially in a novel environment, we want to learn quickly. Since MC is unbiased by the initial estimate of the SR, it would theoretically be beneficial to have more replays in an unfamiliar environment. When the environment is familiar, we want low variance, and TD learning would theoretically be beneficial. We confirm this logic using our spiking neural networks, and show how we can have both quick learning and low error at convergence if we proportionally have more replays at the first trials in a novel environment (Figure 4a-e). In contrast, when having an equal proportion of replays throughout the whole simulation, we do not yield as quick learning as MC and as low asymptotic error as TD (Supplementary Figure 3). Interestingly, the pattern of proportionally more replays in novel environments versus familiar environments has also been experimentally observed [Cheng and Frank, 2008] (Figure 4f).

### 2.6 Leveraging replays to learn novel trajectories

In the previous section, the replays re-activated the same trajectories as seen during behaviour. In this section, we extend this idea and show how in our model replays can be useful during learning even when the re-activated trajectories were not directly experienced during behaviour.

For this purpose, we reproduce an place-avoidance experiment from [Wu et al., 2017]. In short, rats are allowed to freely explore a linear track on day 1. Half of the track is dark, while the other half is bright. On day 2, the animals did four trials separated by resting periods: in the first trial (pre), the animals were free to explore the track; in the second trial (shock), they started in the light zone and received two mild footshocks when entering in the shock zone; in the third and fourth trial (post and re-exposure respectively) they were allowed to freely explore the track again, but starting from the light zone or the shock zone respectively (Figure 5a). In the study, it was reported that during the post trial, animals tended to stay in the light zone and forward replays from the current position to the shock zone were observed when the animals reached the boundary between the light and the dark zone (Figure 5b,c).

**Figure 5.**
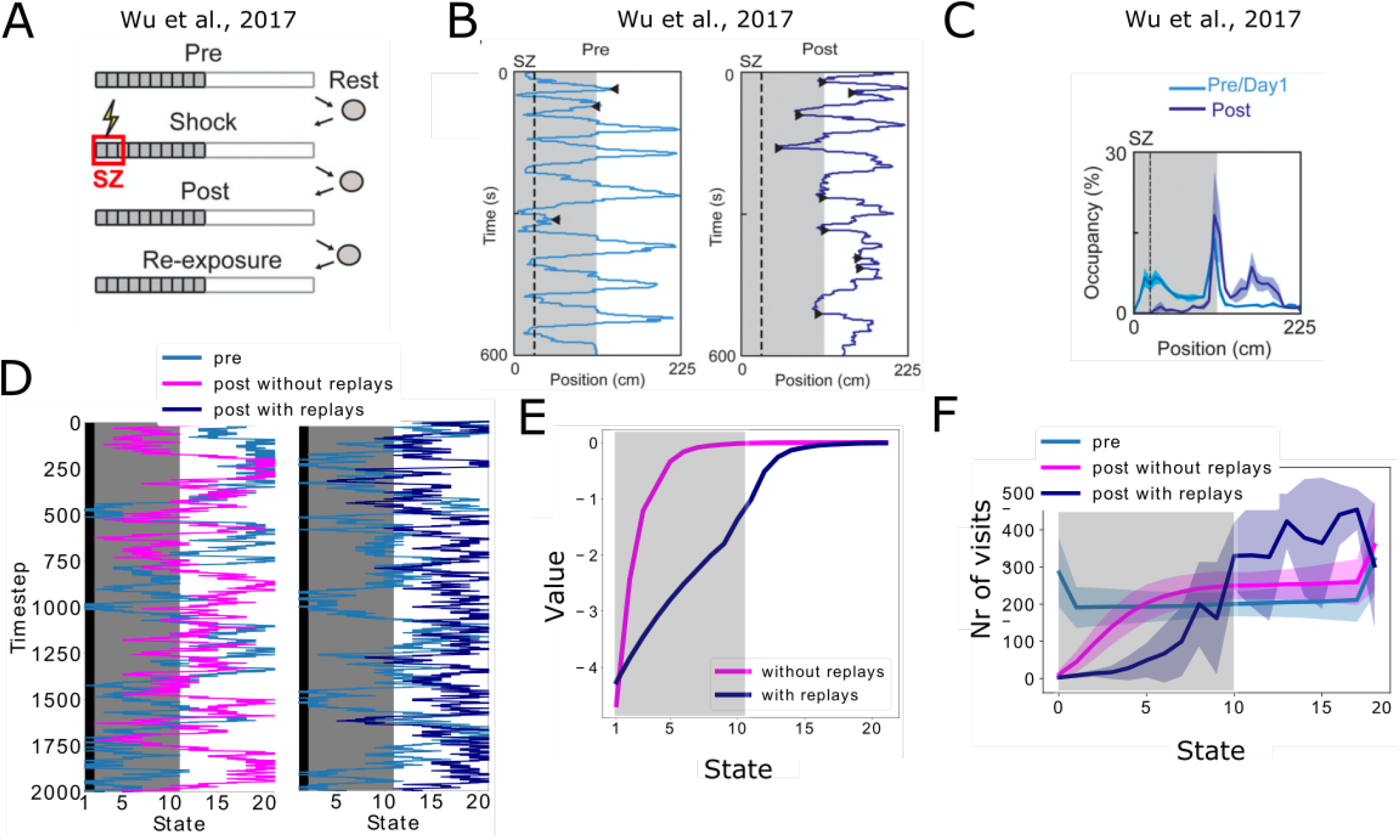
(A-C) Data from Wu et al. [2017]. (A) Experimental protocol: the animal is first allowed to freely run in the track (Pre). In the next trial, a footshock is given in the shock zone (Shock). In subsequent trials the animal is again free to run in the track (Post, Re-exposure). Figure redrawn from Wu et al. [2017]. (B) In the Post trial, the animal learned to avoid the shock zone completely and the also mostly avoids the dark area of the track. Figure redrawn from Wu et al. [2017]. (C) Time spent per location confirms that the animal prefers the light part of the track in the Post trial. Figure redrawn from Wu et al. [2017]. (D) Mimicking the results from Wu et al. [2017], the shock zone is indicated by the black region, the dark zone by the grey region and the light zone by the white region. Left: without replays, the agent keeps extensively exploring the dark zone even after having experienced the shock. Right: with replays, the agent largely avoids entering the dark zone after having experienced the shock (replays not shown). (E) The value of each state in the cases with and without replays. (F) Occupancy of each state in our simulations and for the various trials. Solid line and shaded areas denote the average and standard deviation over 100 simulations respectively. Notice we do not reproduce the peak of occupancy at the middle of the track as seen in panel c, since our simplified model assumes the same amount of time is spent in each state.

We simulated a simplified version of this task. Our simulated agent moves through the linear track following a softmax policy, and all states have equal value during the first phase (pre) (Figure 5d, blue trajectories). Then, the agent is allowed to move through the linear track until it reaches the shock zone and experiences a negative reward. Finally, the third phase is similar as the first phase and the animal is free to explore the track. Two versions of this third phase were simulated. In one version, there are no replays (Figure 5d, orange trajectories in left panel), while in the second version a forward replay until the shock zone is simulated every time the agent enters the middle state (Figure 5d, orange trajectories in right panel, replays not shown). The replays affect the learning of the successor representation and the negative reward information is propagated towards the decision point in the middle of the track. The states in the dark zone therefore have lower value compared to the case without replays (Figure 5e). In turn, this different value affects the policy of the agent which now tends to avoid the dark zone all together, while the agent without replays still occupies many states of the dark zone as much as states in the light zone (Figure 5f). Moreover, even when doubling the amount of SR updates in the scenario without replays, the behaviour of the agent remains unaltered (Supplementary Figure 4). This shows that it is not the amount of updates, but the type of policy that is important when updating the SR, and how using a different policy in the replay activity can significantly alter behaviour.

Our setup for this simulation is simplified, and does not aim to reproduce the complex decision making of the rats. Observe for example the peak of occupancy of the middle state by the animals (Figure 5c), which is not captured by our model because we assume the agent to spend the same amount of time in each state. Nonetheless, it is interesting to see how replaying trajectories that were not directly experienced before, in combination with a model allowing replays to affect the learning of a cognitive map, can substantially influence the final policy of an agent and the overall performance. This mental imagination of trajectories could be exploited to refine our cognitive maps, avoiding unfavourable locations or finding shortcuts to rewards.

## 3. Discussion

In this article, we investigated how a spiking neural network model of the hippocampus can learn the successor representation. Interestingly, we show that the updates in synaptic weights resulting from our biologically plausible STDP rule are equivalent to TD(*λ*) updates, a well-known and powerful reinforcement learning algorithm.

For the neuroscience field, uncovering such a connection between STDP and TD(*λ*) shows how, using minimal assumptions, a theoretically grounded learning algorithm can emerge from a biological implementation of plasticity. This framework could be further generalized to areas such as the neocortex. Another interesting link between biologically plausible learning and TD(*λ*) has been shown by [Brea et al., 2016] in the context of a 2-compartment single neuron. In our network model however, we can relate the *λ* parameter with the type of neuronal activity (rate versus temporal), and the *γ* and *η* parameters with biological parameters such as firing rate, the time the agent spends in a state and plasticity amplitudes.

Our knowledge of the TD(*λ*) algorithm allowed us to speculate about the role replays can play during learning, as well as providing an explanation on how we can learn relationships over behavioural timescales with synaptic plasticity rules on the order of milliseconds - a property also found in the single neuron model in [Brea et al., 2016]. Finally, our framework unifies the rate versus temporal coding ideas that have long been debated. With our model, we provide insights into how temporally precise activations on the one hand and rate-based activations on the other hand can both contribute to the same learning goal and with the same underlying STDP rules.

Moreover, the STDP rule we put forward here is very similar to rules that have been described before [van Rossum et al., 2012; Waddington et al., 2012; Shouval et al., 2002], but the neural activities are slightly more constrained in our framework. Notably, we required external place-tuned inputs to CA3 and after a little delay to CA1. Such inputs could be provided by the entorhinal cortex, while the timing could be regulated by the theta-oscillations known to be present during behaviour. In our framework, also the replays need to activate CA1 place cells after a little delay with respect to CA3 place cells. It is not known if replays in feed-forward layers obey such a timing, and it would therefore be interesting to design experiments investigating the relation between replays in subsequent layers.

Multiple ideas from reinforcement learning, such as TD(*λ*), state-dependent discounting and the successor representation, emerge quite naturally from our simple biologically plausible setting. We propose here that time and space can be discounted differently, as well as that combining experience replay and behavioural learning can reduce the bias and variance. The flexibility to change the discounting factor by modulating firing rates and plasticity parameters — which is ubiquitous in biological neural networks — suggests that the brain could very flexibly encode information in a cognitive map. Finally, our framework does not need an explicit encoding of the prediction error. Instead, the synaptic weights encode the predictions of future state occupancies, and the prediction error is encoded implicitly by combining these synaptic weights and the actually observed states in the STDP rule.

## 4. Methods

### 4.0 The Successor Representation

In a tabular environment, we define the value of a state *s* as being the expected cumulative reward that an agent will receive following a certain policy starting in *s*. The future rewards are multiplied by a factor 0 < *γ*^*n*^ ≤ 1, where *n* is the number of steps until reaching the reward location and 0 < *γ* ≤ 1 is the delay discount factor. It is usual to use 0 < *γ* < 1, which ensures earlier rewards are given more importance compared to later rewards. Formally, the value of a state *s* under a certain policy *π* is defined as

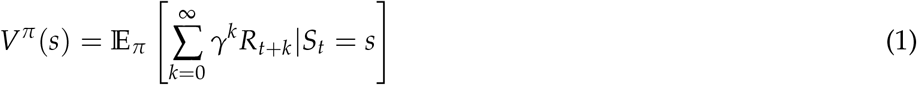

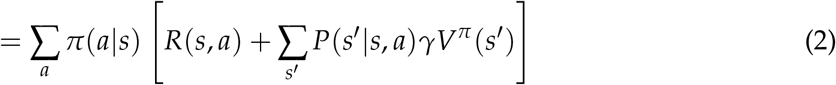

Here, *a* denote the actions, *R*(*s, a*) is the reward function and *P*(*s*′ |*s, a*) is the transition function, i.e. the probability that taking an action *a* in state *s* will result in a transition to state *s*′. Following [Dayan, 1993], we can decompose the value function into the inner product of reward function and successor matrix

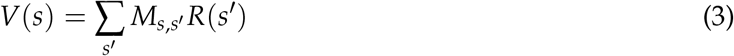

with

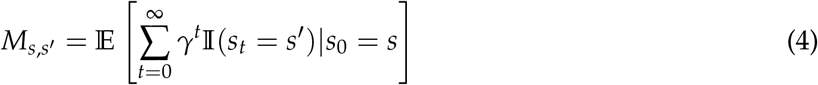

This representation is known as the successor representation (SR), where each element *M*_*ij*_ represents the expected future occupancy of state *j* when in state *i*. By decomposing the value into the SR and the reward function (equation 3), relearning the state values *V* after changing the reward function is fast, similar to model-based learning. At the same time, the SR can be learned in a model-free manner, using for example temporal difference (TD) learning [Russek et al., 2017].

### 4.0 Derivation of the TD(*λ*) update for the SR

The TD(*λ*) update for the SR is then implemented according to (see e.g. [Sutton and Barto, 1998])

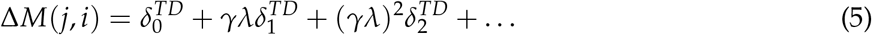

Using 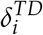 for the TD error at step *i* and *δ*_*xy*_ for the Kronecker delta,

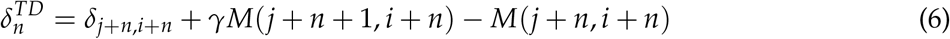

corresponds to the TD error for element *M*(*j* + *n, i* + *n*) of the successor representation after the transition from state *j* + *n* to state *j* + *n* + 1. Combining equations 5 and 6, we find

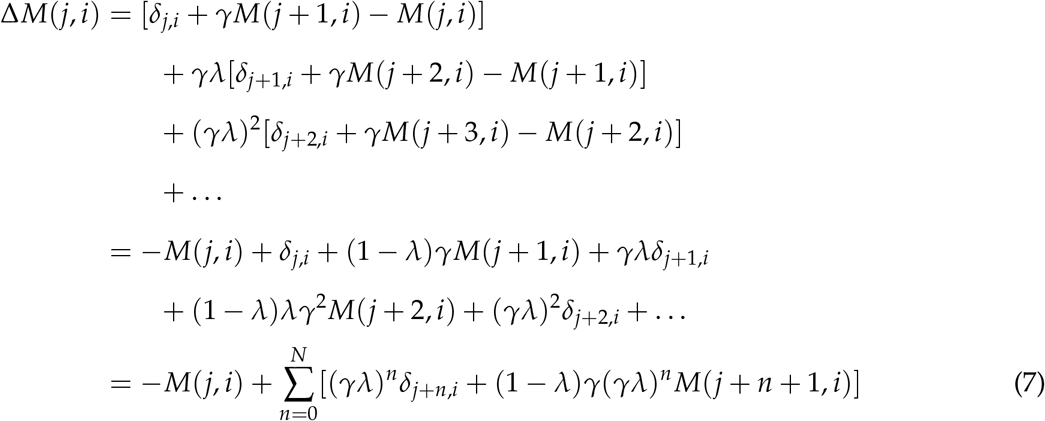

and

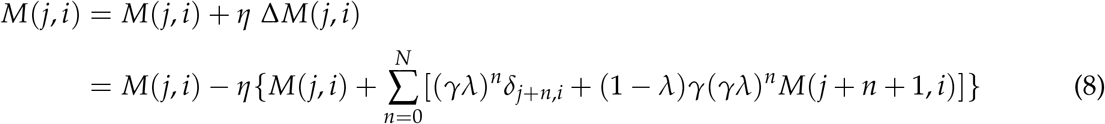

### 4.0 Neural network model

#### Plasticity rule

The synaptic plasticity rule (Figure 1d) consists of a weight-dependent depression for presynaptic spikes and a spike-timing dependent potentiation, given by

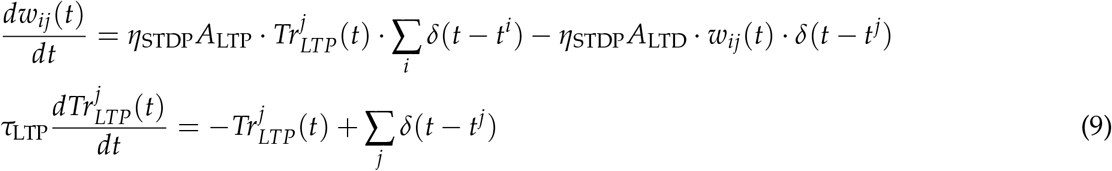

Here, *w*_*ij*_ represents the synaptic connection from presynaptic neuron *j* to postsynaptic neuron *i*, 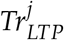 is the plasticity trace, a low-pass filter of the presynaptic spiketrain with timeconstant *τ*_LTP_, *t*^*j*^ and *t*^*i*^ are the spike times of the postsynaptic and presynaptic neuron respectively, *A*_LTP_ and *A*_LTD_ are the amplitudes of potentiation and depression respectively, *η*_STDP_ is the learning rate for STDP and the *δ*(·) denotes the Dirac delta function.

#### Place cell activation

We assume that each state in the environment is represented by a population of place cells in the network. In our model, this is achieved by delivering place-tuned currents to the neurons. Whenever a state *S* = *j* is entered, the presynaptic neurons encoding state *j* start firing at a constant rate *ρ*^*pre*^ for a time *θ*, following a Poisson process with parameter 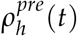. The other presynaptic neurons are assumed to be silent:

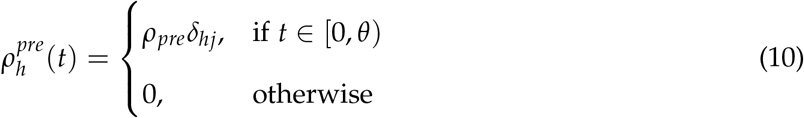

where the Kronecker delta function is defined as *δ*_*hj*_ = 1 if *h* = *j* and zero otherwise. Here we use the index *j* to denote any neuron belonging to the population of neurons encoding state *j*. After a short delay, at time *t*\*, a similar current *ρ*^*bias*^ is delivered to the postsynaptic neuron encoding state *j*, for a duration of time *ω*.

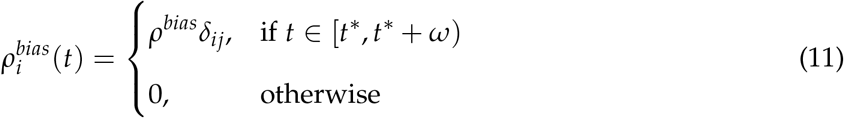

Besides the place-tuned input current, CA1 neurons receive inputs from the presynaptic layer (CA3). The postsynaptic potential 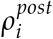 when the agent is in state j is thus given by

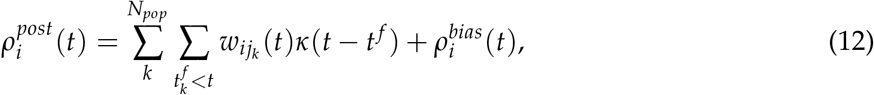

with the first sum running over all *N*_*pop*_ presynaptic neurons encoding state *j*, and the second sum over all presynaptic firing times 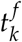 of neuron *k* happened before *t*. The excitatory postsynaptic current *κ* is modelled as an exponential decay described as 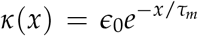 for *x* ≥ 0 and zero otherwise. Each CA1 neuron *i* fires following an inhomogeneous Poisson process with rate 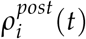.

Note that, in most simulations we will use a single neuron in the population *N*_*pop*_ = 1. In addition, we normally set *t*\* = *θ* and *ω* = *T* − *θ*. However, we will keep these as explicit parameters for theoretical purposes.

#### Equivalence with TD(*λ*)

##### Total plasticity update

Since we have the mathematical equation for the plasticity rule, and the firing rates of CA3 and CA1 neurons follow a Poisson process, we can calculate analytically the average total weight change for the synapse *w*_*ij*_, given a certain trajectory (details in the Supplementary Materials). We find that:

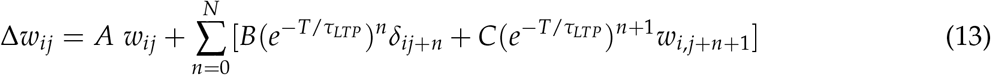

where *N* is the number of states until the end of the trajectory and

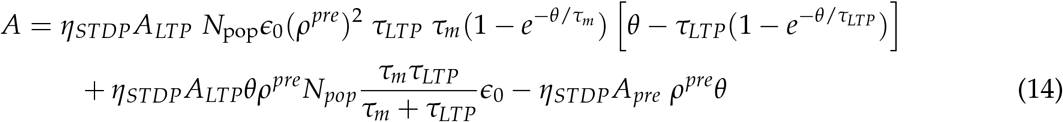

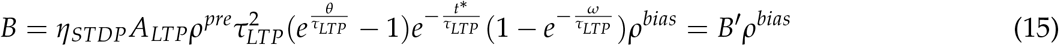

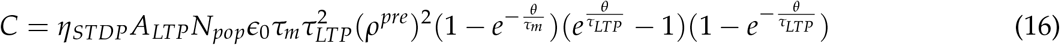

##### Comparison with TD(*λ*)

Comparing the total weight change due to STDP (equation 13) to the TD(*λ*) update (equation 8), we can see that the two equations are very similar in form:

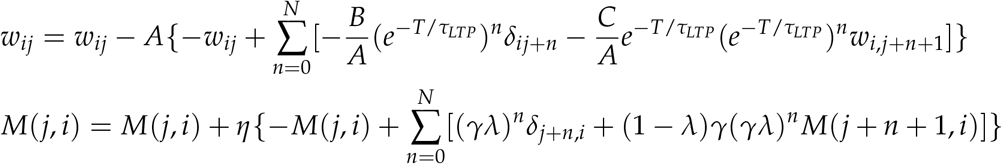

We impose *w*_*ij*_ = *M*(*j, i*), and find:

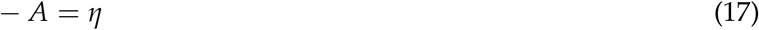

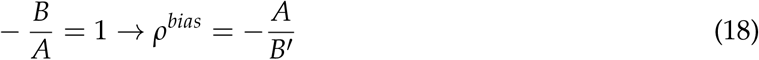

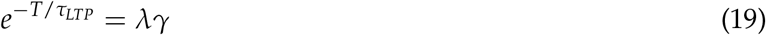

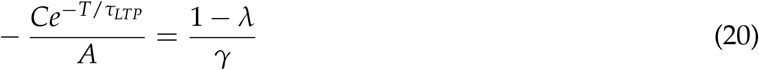

Hence, our plasticity rule is learning the Successor Representation through a TD(*λ*) model with parameters:

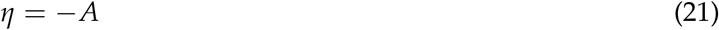

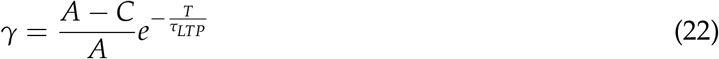

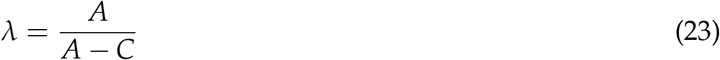

To ensure the learning rate *η* is positive, one condition resulting from equation 21 is

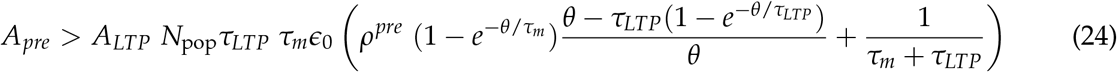

#### Learning during normal behaviour (*θ* >> *τ*_*LTP*_)

During normal behaviour, we assume the place-tuned currents are on larger timescales than the plasticity constants: *θ, ω* >> *τ*_*LTP*_. We can see from equations 14 and 16 that the factor *A* grows linearly with *θ* while *C* grows exponentially with *θ*. From equation 23, we then have

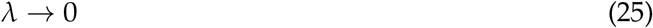

(See also Supplementary Figure 1).

#### Learning during replays (*θ* << *τ*_*LTP*_)

##### Assumptions

For the replay model we assume the place-tuned currents are impulses, which make the neurons emit exactly one spike at a given time. Specifically, we can make the duration of the place-tuned currents go to 0,

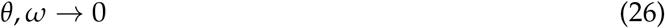

while the intensity of the currents goes to infinity. For simplicity, we will take:

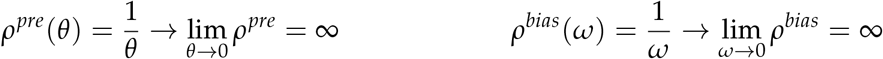

Furthermore, we assume that the contribution of the postsynaptic currents due to the single presynaptic spikes is negligible in terms of driving plasticity, allowing us to set

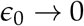

##### Calculations of TD parameters

Given the assumptions above, we can see from equations 14 and 16 that:

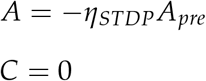

For equation 15, we can use the Taylor expansion for 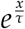 around *x* = 0, such that: 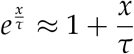:

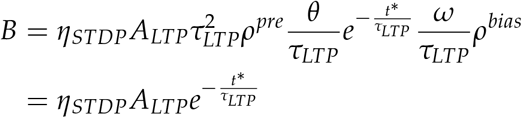

Using equations 21, 22, 23 and 18, we can calculate the parameters and constraints for the TD model:

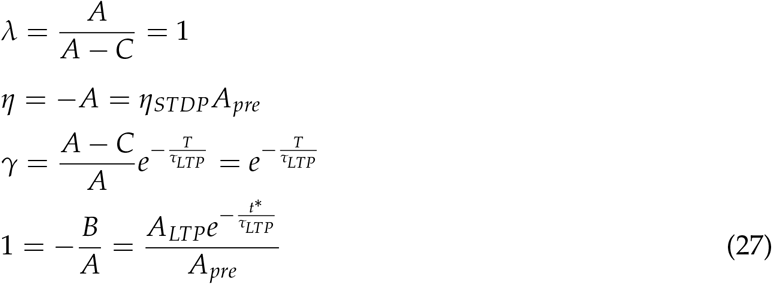

As expected, the bootstrapping parameter *λ* = 1 (see also Supplementary Figure 1).

#### Alternative derivation of replay model

##### Place cell activation

We model a replay event as a precise temporal sequence of spikes. Since every neuron represents a state in the environment, a replay sequence reproduces a trajectory of states. We assume that, when the agent is in state *S* = *j*, the neurons representing state *j* fire *n*_*pre*_ spikes at some point in the time interval *t* ∈ [0, *σ*], where the exact firing times are uniformly sampled. After a short delay, the CA1 neurons representing state *j* fire *n*_*post*_ spikes at a time uniformly sampled from the interval [*t*\*, *t*\* + *σ*]. The time between two consecutive state visits is *T*. The exact number of spikes in each replay event is random but small. Specifically, it is sampled from the set {0, 1, 2} according to the probability vector 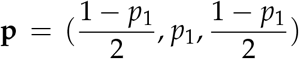. It is worth noting here that other implementations are possible but that we assume the average number of spikes in each state is 1, and that the average time between a presynaptic and a postsynaptic spike is *t*\*. The model could be further generalized for a higher number of average spikes per state.

##### Plasticity update

We can consider again our learning rule, composed of a positive pre-post potentiation window and presynaptic weight-dependent depression (equation 9). Let’s consider the synapse *w*_*ij*_, we can see that on average the total amount of depression will be determined by the number of times the state *j* is visited in the trajectory replayed:

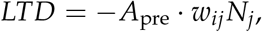

where *N*_*j*_ is the number of times the state *j* is visited. The amount of potentiation will be determined, instead, by the time difference between the postsynaptic and presynaptic firing times, which encode the distance between state *j* and state *i*:

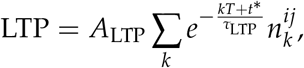

where 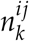 represents the number of times the agent visited state *i* k steps after *j*. Combining the equations above we find that:

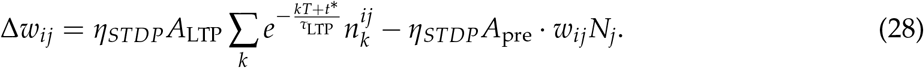

If we assume that the this value has converged to its stationary state, Δ*w*_*ij*_ = 0:

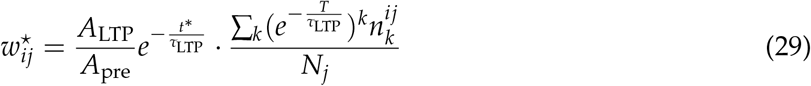

##### Comparison with online Monte Carlo learning

Given the stable weight *w*\* from equation 29, we can impose that:

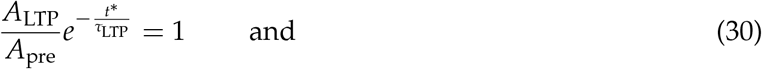

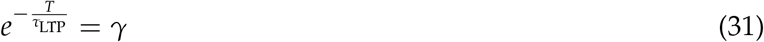

we find that the stable weight is:

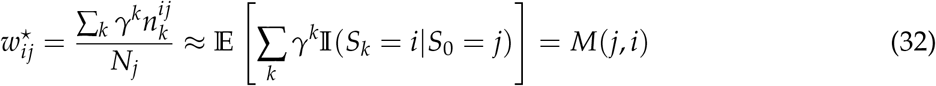

which is the definition of the Successor Representation matrix (equation 4). Indeed, 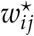 is computing the sample mean of the discounted distance between states *i* and *j*, which is equivalent to performing an every-state Monte Carlo or TD(*λ*=1) update. Notably, from equation 28, we have that the learning rate for the Monte Carlo update is given by:

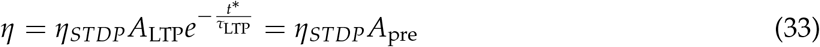

### 4.0 Simulation details for Figure 2

A linear track with 4 states is simulated. The policy of the agent in this simulation is to traverse the track from left to right, with one epoch consisting of starting in state 1 and ending in state 4. One simulation consists of 50 epochs, and we re-run the whole simulation ten times with different random seeds. Over these ten seeds, mean and standard deviation of the synaptic weights are recorded after every epoch.

Our neural network consists of two layers, each with a single neuron per state (as in Figure 1). Synaptic connections are made from each presynaptic neuron to all postsynaptic neurons, resulting in a 4-by-4 matrix which is initialized as the identity matrix. The plasticity rule and neuronal activations follow equations 9 to 12.

The STDP parameters are listed in Table 1.

**Table 1.** Parameters used for the spiking network.

|  |  |
| --- | --- |
| $\epsilon_0$ | 1 |
| $\rho_{\text{pre}}$ | $0.1 \text{ ms}^{-1}$ |
| $\tau_m$ | 2 ms |
| $N_{\text{post}}$ | 1 |
| $N_{\text{pre\_tot}}$ | 1 |
| $N_{\text{pre}}$ | 1 |
| stepsize | 0.01 ms |
| $\eta_{\text{stdp}}$ | 0.003 |
| $\tau_{\text{LTP}}$ | 60 ms |
| $A_{\text{LTP}}$ | $1 \text{ ms}^{-1}$ |

To obey equation 24, we set *A*_pre_ equal to the right hand side augmented with 5.

For the behavioural case, we choose T=100 ms, *θ*=80 ms, *ω* = *T* − *θ*, which correspond to TD(*λ*) parameters *λ* = 0.21, *γ* = 0.89, *η* = 0.12.

In the replay case, we have a sequence of single spike per neuron (see Figure 2b and section 4). Following equation 27, we choose *T* = − log (*γ*) *τ*_*LTP*_ ≈ 7ms, where *γ* and *τ*_*LTP*_ are the same as in table 1. We set *θ* = 2ms and *σ* = 0.5ms. By setting the 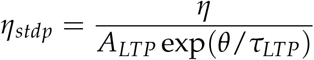, the corresponding TD(*λ*) parameters are *λ* = 1, *γ* = 0.89, *η* = 0.12 just as in the behavioural case.

Finally, we choose *p*_1_ = 0.15, in order to achieve an equal amount of noise due to the random spiking as in the case of behavioural activity (see Supplementary Figure 2).

### 4.0 Simulation details for Figure 3

Using the same neural network and plasticity parameters as the behavioural learning in Figure 2 (see previous section), we simulate the linear track in the following two situations:

- The third state has T = 200 ms instead of 100 ms. All other parameters remain the same as in Figure 2. Results plotted in Figure 3E.
- The third state has *ρ*_pre_ = 0.2 ms^−1^ instead of 0.1 ms^−1^. All other parameters remain the same as in Figure 2. Results plotted in Figure 3F.

### 4.0 Simulation details for Figure 4

A linear track with 3 states is simulated, and the agent has 50% probability to move left or right in each state (see Figure 4A). One epoch lasts until the agent reaches one of the *STOP* locations.

We then use the same neural network and plasticity parameters as used for Figure 2. We simulate three scenarios:

- Only replay-based learning during all epochs (no behavioural learning). This scenario corresponds to *MC STDP* in Figure 4B and to 4C.
- Mixed learning using both behaviour and replays. The probability for an epoch to be a replay is decaying over time following exp(−*i*/6), with *i* the epoch number. This scenario corresponds to *Mix STDP* in Figure 4B and to 4E.
- Only behavioural learning during all epochs (no replays). This scenario corresponds to *TD STDP* in Figure 4B and to 4D.

### 4.0 Simulation details for Figure 5

A linear track with 21 states is simulated. The SR is initialized as the identity matrix, and the reward vector (containing the reward at each state) is also initialized as the zero vector. We simulate the learning of the SR during behaviour using the theoretical TD(0) updates and during replays using the theoretical TD(1) updates. The value of each state is then calculated as the matrix-vector product between the SR and the reward vector, resulting in an initial value of zero for each state.

The policy of the agent is a softmax policy (i.e. the probability to move to neighbouring states is equal to the softmax of the values of those neighbouring states). The first time the agent reaches the leftmost state of the track (state 1), the negative reward of -2 is revealed, mimicking the shock in the actual experiments, and the reward vector is updated accordingly for this state.

We now simulate two scenarios: in the first scenario, the agent always follows the softmax policy and no replays are triggered (see Figure 5D, left panel). In the second scenario, every time the agent enters the dark zone from the light zone (i.e. transitions from state 12 to state 11 in our simulation), a replay is triggered from that state until the leftmost state (state 1) (see Figure 5D, right panel). Both scenarios are simulated for 2000 state transitions. We then run these two scenarios 100 times and calculate mean and standard deviation of state occupancies (Figure 5F).

Finally, since the second scenario has more SR updates than the first scenario, we also simulate the first scenario for 4000 state transitions (Supplementary Figure 4) and show how the observed behaviour of Figure 5 is unaffected by this.

## 6. Supplementary materials

### 6.1 Analytical derivations for the total weight change in the behavioural model

#### Presynaptic rate during state *j*

Whenever a state *S* = *j* is entered, the presynaptic neurons encoding state *j* start firing at a constant rate *ρ*^*pre*^ for a time *θ*, following a Poisson process with parameter 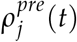:

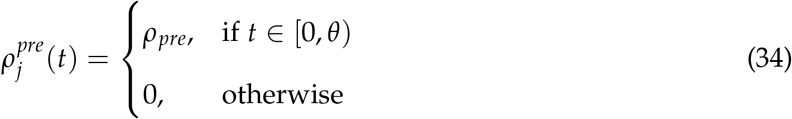

The other presynaptic neurons are silent.

#### Postsynaptic rate during state *j*

The average postsynaptic rate can be calculated as follows. The probability of a presynaptic spike between *t* and *t* + *dt* is equal to *ρ*_*j*_(*t*)*dt*. The size of the presynaptic population encoding state *j* is equal to *N*_pop_ and each excitatory postsynaptic potential (EPSP) is modelled by an immediate jump with amplitude *ϵ*_0_*w*_*ij*_, followed by exponential decay of EPSP with time constant *τ*_*m*_.

Following equation 12 in the main paper, reproduced below,

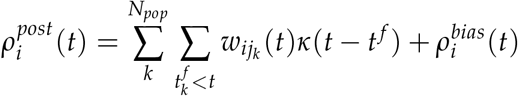

we find that the average postsynaptic potential at time *t* is given by (assuming t=0 when entering the state *j*):

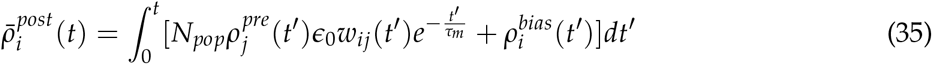

We assume that *w*_*ij*_(*t*) changes slowly compared to the timescale *θ* allowing us to consider the weight constant during that time. We can then approximate the average postsynaptic rate as:

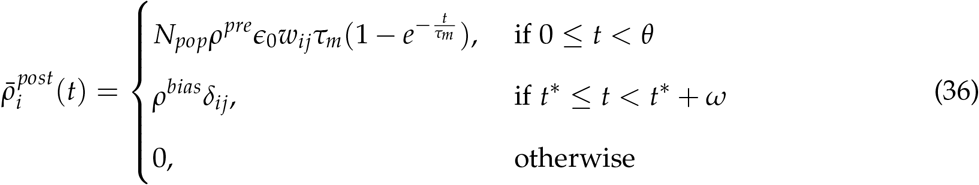

If *t*^⋆^ < *θ*, both the first and the second term will contribute to the postsynaptic rate in the time between *t*^⋆^ and *θ*.

#### LTP trace during state *j*

Given equation 9 in the main paper, reproduced below,

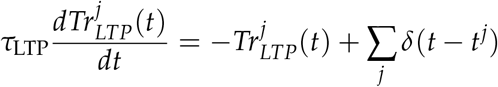

and combined with equation 34, we can calculate the evolution of the LTP trace for neuron *j* during state *j*:

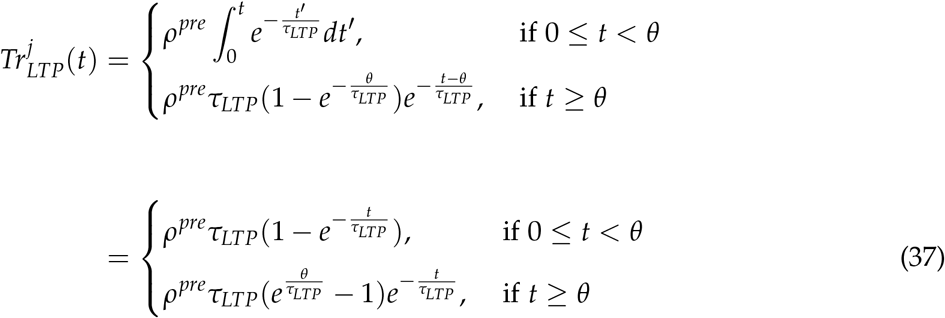

For 0 ≤ *t* < *θ*, the presynaptic neuron *j* is active and therefore the trace builds up with the presynaptic spikes, for *t* ≥ *θ*, the trace decays exponentially with time constant *τ*_*LTP*_.

#### Total amount of LTP during state *j*

Following [Kempter et al., 1999], first we calculate the amount of LTP without taking into account spike-to-spike correlation:

The probability for a postsynaptic spike between *t* and *t* + *dt* is 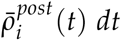. The amount of LTP due to a single spike at time *t* is 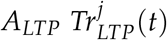. Hence, combining equations 36 and 37, the total amount of LTP during a state (i.e. between time 0 and *T*) becomes:

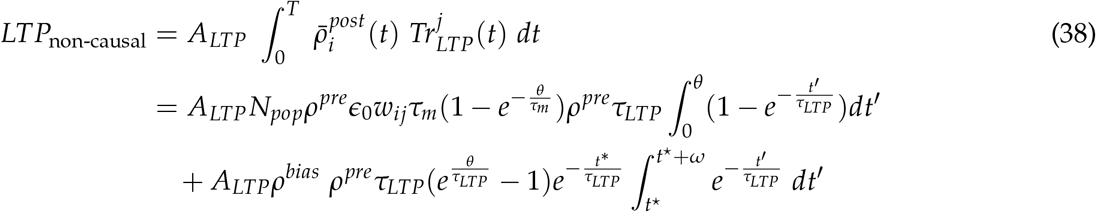

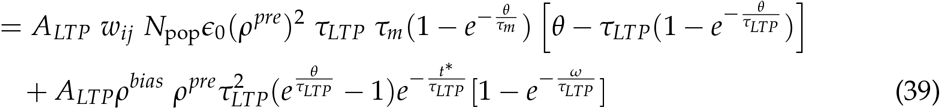

Following [Kempter et al., 1999], the amount of LTP due to the causal part (each presynaptic spike temporarily increase the probability of a postsynaptic spike) is given by:

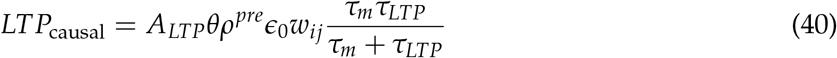

Combining equations for the non-causal 39 and causal 40 parts, we get the *total* amount of LTP during a state (assuming *τ*_*m*_ << *τ*_*LTP*_):

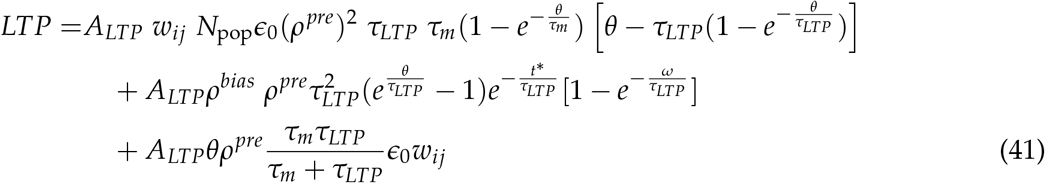

#### Total amount of LTD during state *j*

There is a weight-dependent depression for each presynaptic spike, hence the amount of LTD during a state is given by:

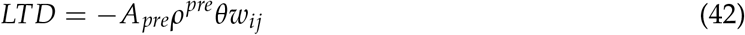

#### Total plasticity during state *j*

Combining equations 41 and 42, we can calculate the total amount of plasticity during the time the agent spends in the current state *j*:

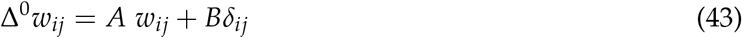

with

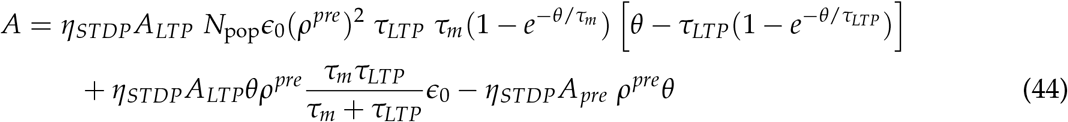

and

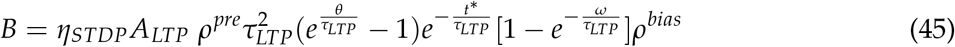

#### Plasticity due to states transitioning

Once the agent leaves state *j*, the decaying LTP trace can still cause potentiation due to the activity in the following states, *j* + *n*, with *n* = 1, 2,…. Given that the agent spends a time *T* in each state, we find that the agent visits state *j* + *n* during time *t* ∈ [*nT, nT* + *T*). We will now calculate the contribution to plasticity due to these state transitions.

#### Postsynaptic rate during the new state *j* + *n*

During state *j* + *n*, the activity of the postsynaptic neurons is driven by the presynaptic neurons coding for *j* + *n*, and the bias current. We can thus generalize equation 36 and find that the average postsynaptic rate 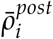 during state *j* + *n* is:

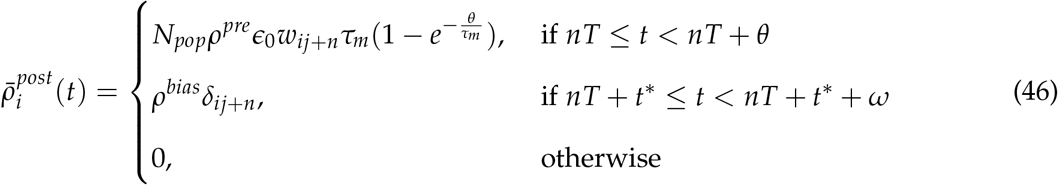

#### LTP trace from state *j*, during the new state *j* + *n*

Following equation 37, we find that the amplitude of the LTP trace from state *j* during state *j* + *n* is:

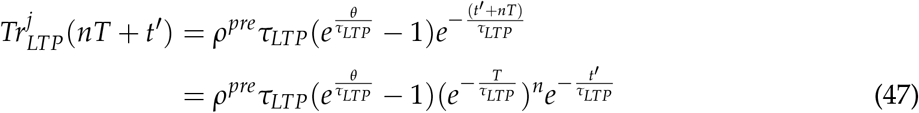

with 0 < *t*′ < *T*

#### LTP due to state transitioning

We can then calculate the amount of LTP between the presynaptic neuron *j* and the postsynaptic neuron *i*, when the agent is in state *j* + *n*. We refer to equation 38 and find:

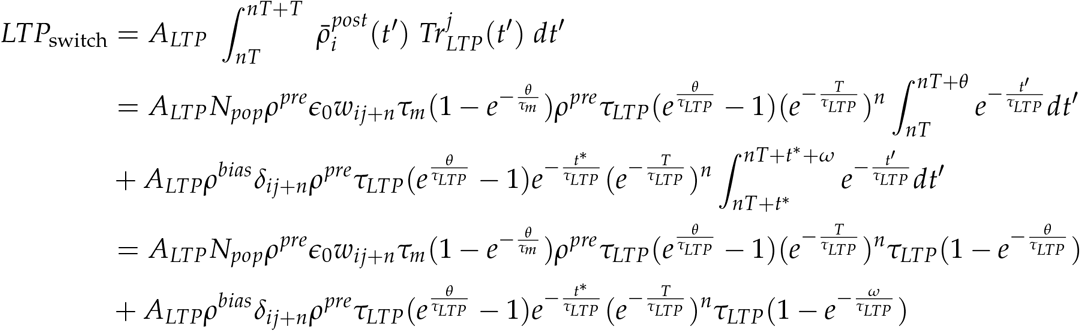

The amount of plasticity in state *j* + *n* when starting from state *j* is thus:

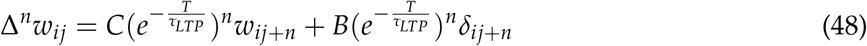

where

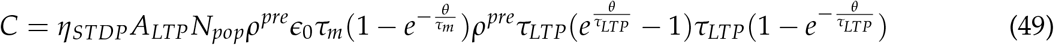

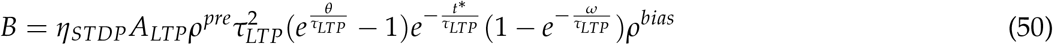

It is worth noting that the parameter B derived here is the same as equation 45.

### 6.1 Summary: total STDP update

If we combine together equations 43 and 48, we have that the total weight change for the synapse *w*_*ij*_ is given by:

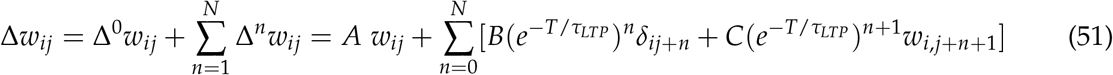

where *N* is the number of states until the end of the trajectory and A, B, C are as defined in equations 44, 45 and 49 respectively.

## 6. Analytical calculations for hyperbolic discounting

From equation 22 in the main paper, we have that, in the behavioural model 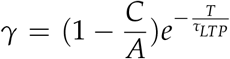.

Here, we will derive an approximation to this value.

If we assume that *θ >> τ*_*m*_, *τ*_*LTP*_, we can approximate *A* and *C* as:

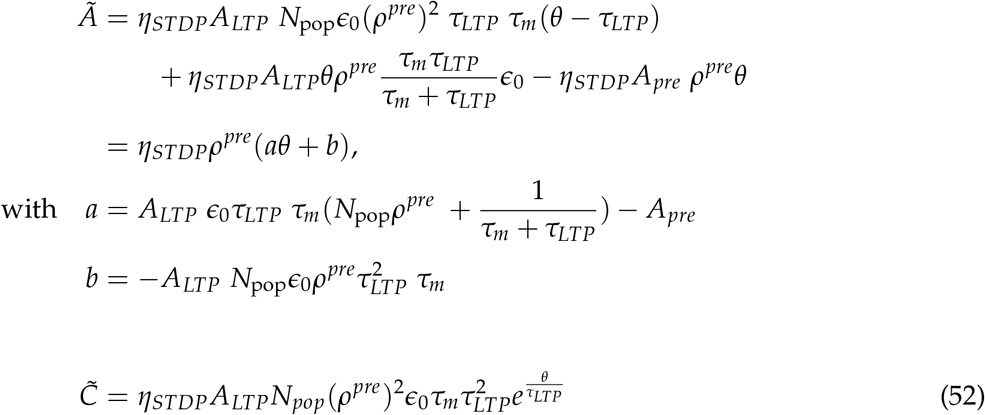

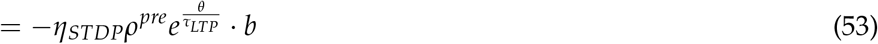

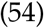

If we define *ψ* such that *θ* + *ψ* = *T*, we can rewrite and approximate the discount parameter as:

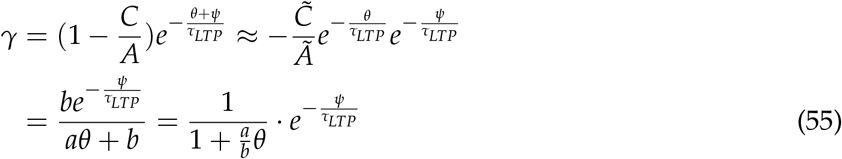

From equation 55, we can see that the discount *γ* follows a hyperbolic function if we increase the duration of the presynaptic current *θ*. If, instead, we vary *ψ*, the discount becomes exponential (Figure 1 a and b).

Notice that this analysis extends to the replay model. Following what was done after equation 26, we can connect the behavioural model with the replay model by making *θ, ϵ*_0_ → 0, which implies *ψ* → *T*. From equation 55 we find that:

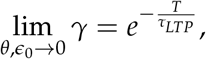

which is exactly the definition of *γ* in the replay model (equations 27 in Methods). For replays, the discount is therefore strictly exponential.

Furthermore, using the same calculations and equations 21 and 19 in the main paper, we can find approximated values for the other parameters too (Supplementary Figure 1 c and d).

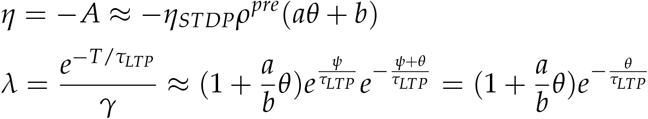

**Supplementary Figure 1.**
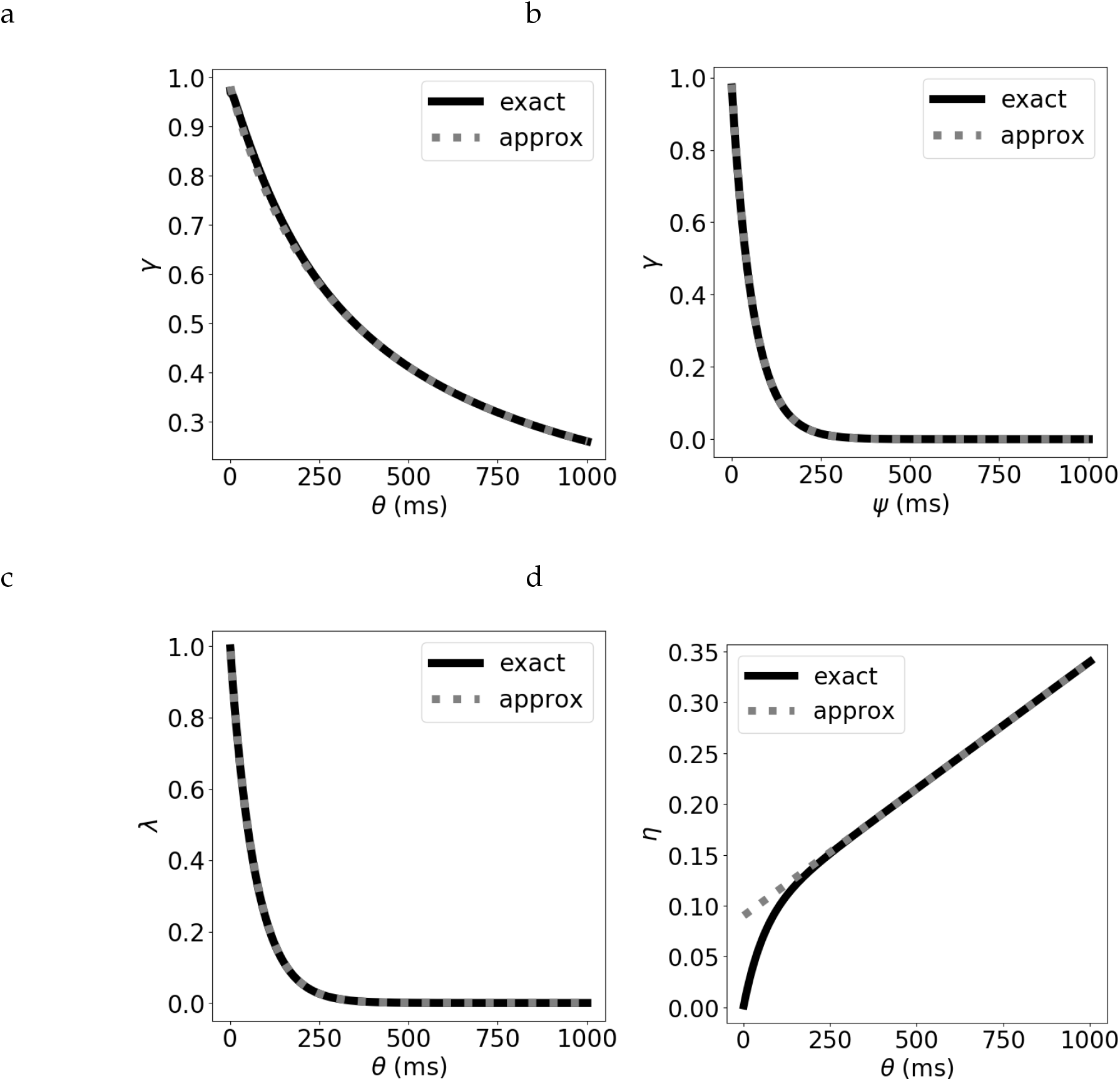
Comparison of the exact and approximate equations for the parameters. The discount parameter *γ* when we vary *T* by: (a) increasing the duration of the presynaptic current *θ*, leading to a hyperbolic discount, and b) increasing *ψ*, while keeping *θ* fixed, leading to an exponential discount. (c) The bootstrapping parameter *λ* for varying *θ*. (d) The learning rate *η* for varying *θ*.

**Supplementary Figure 2.**
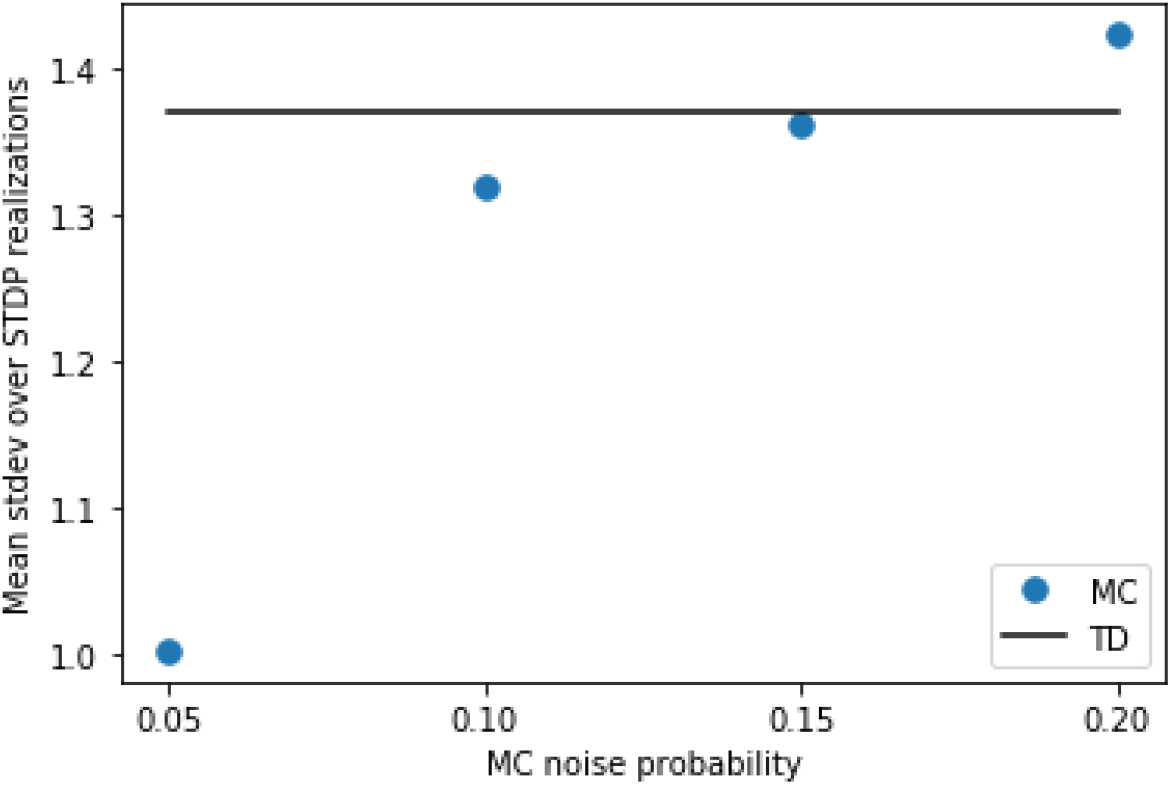
Setting the noise for replays. Using the settings of Figure 4, we calculate the variance caused by random spiking for the behavioural model first. For this purpose, we simulate the same trajectory of the agent 25 times with different random instances of spikes generated from the Poisson process. At each state transition, we calculate the standard deviation of the synaptic weights over the 25 runs. Finally, we calculate the mean of these standard deviations over all states of the trajectory. We find a value of this mean standard deviation just below 1.4 (full line). Then, we try various levels of noise for the replay model, and compute the same mean standard deviation. We find that a probability *p*_1_ = 0.15 yields a similar level of variance due to random spiking as in the behavioural model.

**Supplementary Figure 3.**
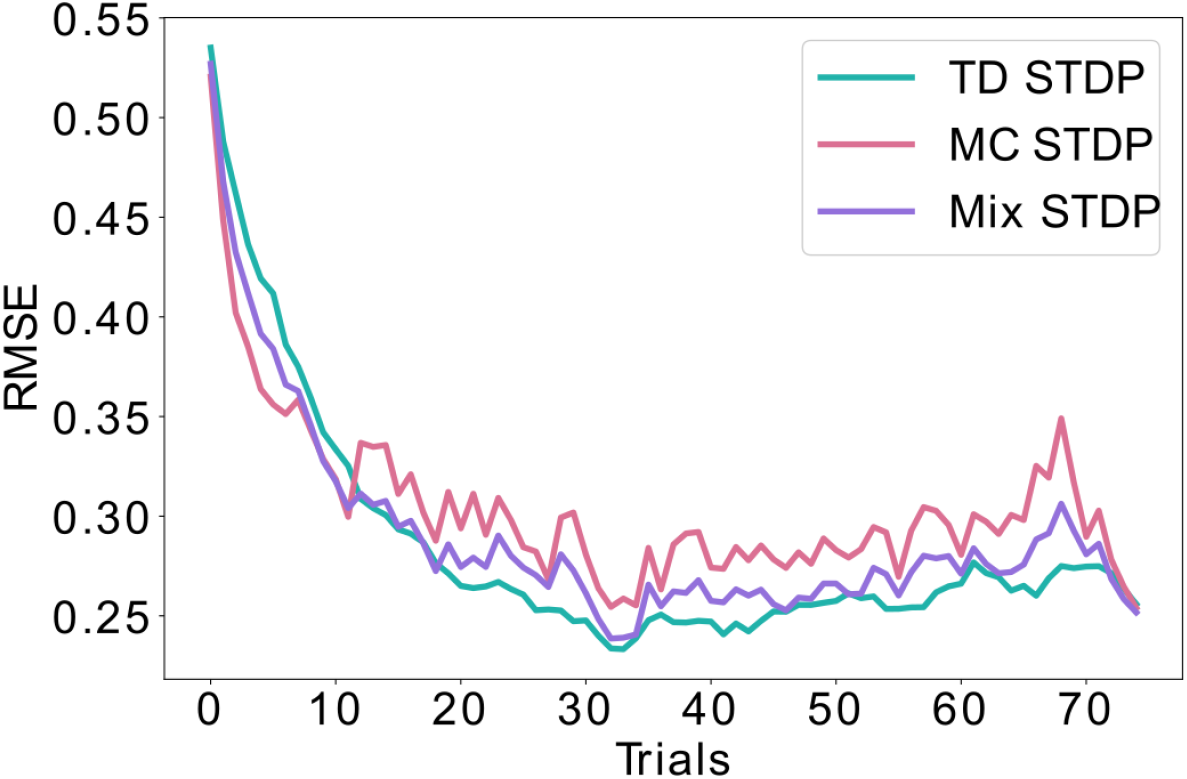
Combining equal amounts of replays and behavioral learning. Unlike figure 4, where the likelihood of replays is exponentially decaying over time, here we simulate equal probability for learning using replays (MC) and using behavioral experience (TD) throughout time. This mixed strategy lowers the bias and the variance to some extent, but lays between the pure MC and pure TD learning, never achieving the minimal error possible. This is in contrast with the exponential decay of replays shown in figure 4, which achieves a minimal error both during early learning (similar to MC) as well as asymptotically (similar to TD)

**Supplementary Figure 4.**
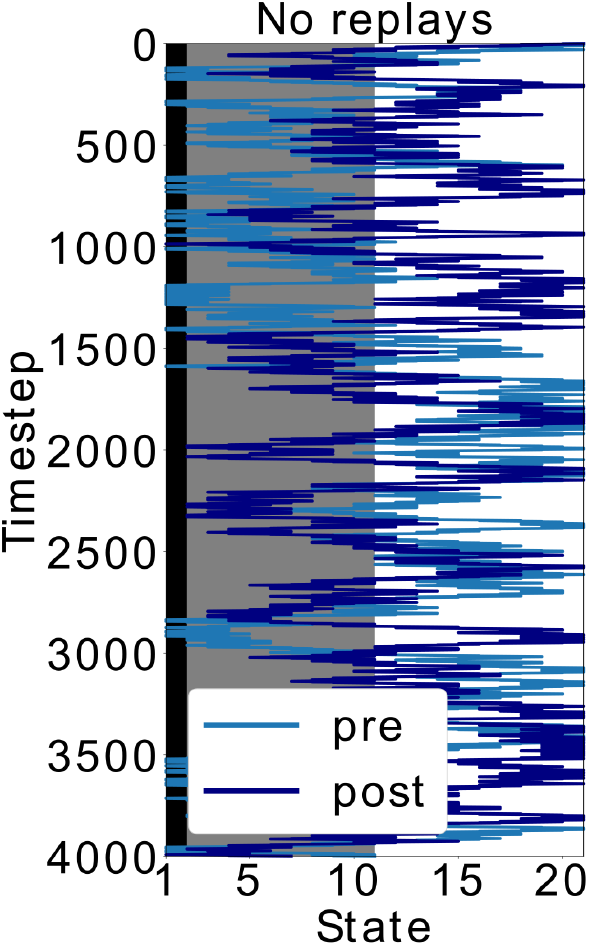
Doubling the time-steps in the scenario without replays. In figure 5, the policy without replays has less SR updates. Here, we simulated this policy but doubled the time (and SR updates). Even with this modification, the agent keeps exploring the dark zone of the track, which shows that it is indeed the type of policy and not the amount of updates that leads to the different behaviour. More specifically, sequential activations of states from the decision point until the shock zone, such as in replays, are different than the softmax policy during behaviour. It is exactly by exploiting this different replay policy that the agent updates the SR differently and avoids the dark zone.

## Notes

### Competing Interest Statement

The authors have declared no competing interest.

